# Visuo-proprioceptive integration and recalibration with multiple visual stimuli

**DOI:** 10.1101/2021.05.12.443755

**Authors:** Nienke B Debats, Herbert Heuer, Christoph Kayser

**Author notes:** CORRESPONDING AUTHOR, Nienke B. Debats, Universitätsstrasse 25, 33615 Bielefeld, Germany.

## Abstract

To organize the plethora of sensory signals from our environment into a coherent percept, our brain relies on the processes of multisensory integration and sensory recalibration. We here asked how visuo-proprioceptive integration and recalibration are shaped by the presence of more than one potentially relevant visual stimulus, hence paving the way to studying multisensory perception under more naturalistic settings with multiple signals per sensory modality. By manipulating the spatio-temporal correspondence between the hand position and two visual stimuli during a cursor-control task, we contrasted two alternative accounts: that only the temporally more proximal signal enters integration and recalibration similar to a winner-takes-all process, or that the influences of both visual signals superpose. Our results show that integration - and likely also recalibration - are shaped by the superposed contributions of multiple stimuli rather than by only individual ones.

## INTRODUCTION

A key aspect of perception is the combination of redundant multisensory signals, that is, of signals received by separate modalities but relating to the same property of an object. Critically, multisensory signals may be discrepant, for example due to sensory noise or environmental distortions such as an echo. Our brain exploits two processes to deal with such discrepancies: multisensory integration and sensory recalibration. Through multisensory integration unisensory signals are combined into a weighted representation, which is typically revealed in studies investigating bimodal judgments (e.g., Ernst and Banks 2002; Ernst 2006; Körding et al. 2007; Cheng et al. 2007; Shams and Beierholm 2010; van Dam et al. 2014; Kayser and Shams 2015; Colonius and Diederich 2020). Sensory recalibration in contrast helps to align unisensory estimates with each other. It is revealed as a unisensory aftereffect – a perceptual bias that occurs after (prolonged) exposure to bimodal discrepancies (e.g., Recanzone 1998; Redding et al. 2005; Henriques and Cressman 2012; Chen and Vroomen 2013; Burr 2014). While typical laboratory tasks usually focus on two isolated unisensory stimuli that are combined during integration or recalibration, real world settings feature an abundance of signals within each modality. This raises the question of how multisensory integration and recalibration operate when multiple apparently redundant signals per modality are present.

To address this question, we investigated hand movements in a cursor-control task, which has previously served as a versatile paradigm to study multisensory integration (e.g., Rand and Heuer 2013; Debats et al. 2017a, b; Debats and Heuer 2018a) and recalibration (e.g., Synofzik et al. 2008; Henriques and Cressman 2012; Rand and Heuer 2019). We asked how perceptual estimates of a proprioceptive target (hand position) are biased by the presence of two (rather than one) visual stimuli presented during the movement. Redundant multisensory signals are often characterized by their spatio-temporal proximity, which is a critical feature determining which signals enter multisensory integration and recalibration (Ernst 2007; Odegaard et al. 2017; Tong et al. 2020) (for reviews see e.g., Shams and Beierholm 2010; Talsma et al. 2010; van Dam et al. 2014). We hence manipulated the two visual stimuli so that they featured the same spatial proximity to the proprioceptive stimulus while differing in their temporal proximity. We hypothesized that temporal proximity would modulate the influence of each visual stimulus during integration and recalibration, with the temporally more proximal stimulus having a stronger influence. Specifically, we compared two putative models describing the influence of each visual stimulus: a ‘winner-takes-all principle’, whereby only one of them would influence proprioceptive judgments, and a ‘superposition principle’, whereby the influences of both visual stimuli would be summed.

So far, only few studies have investigated how multiple stimuli per sensory modality contribute to multisensory integration or recalibration. In a study with two visual stimuli flanking a sound, the perceived sound location was biased towards the visual stimulus exhibiting a stronger tendency to integrate when presented in an isolated audio-visual pair (Tong et al. 2020), while in another audiovisual study, directing attention to one of the two visual stimuli seemed to have little influence on integration (Bertelson et al. 2000). A systematic review on the influence of attention in multisensory integration posed that when multiple stimuli are presented in one modality, they may collectively enter multisensory integration as long as their individual saliency is sufficiently high and both relate to the respective cross-modal stimulus via some pattern of spatio-temporal correlation (Talsma et al. 2010). These results suggest that in our cursor-control task the perceived hand position might indeed be biased toward the temporally more proximal stimulus, for which one would expect a stronger influence when presented as the only visual cue. However, it is not self-evident that findings with audio-visual stimuli, for which participants are observers, generalize to visuo-proprioceptive stimuli, which are actively produced by the participants. Among other differences, only for this type of task the discrimination between reafferent and exafferent visual information is required (cf. Reichenbach et al. 2014). Moreover, the available results obtained with multiple visual stimuli do not provide direct insights on the two competing hypotheses of how multiple cues engage in integration or recalibration.

We here report the results of two experiments. In the first, we examined multisensory integration and recalibration while presenting two visual cues with equivalent spatial distance to the hand position but differing in their temporal distance (synchronous vs. delayed). We found that both integration and recalibration biases were dominated by the synchronous visual signal. In the second experiment, we directly contrasted the winner-takes-all and the superposition hypotheses by comparing the integration biases observed in the presence of both visual stimuli with the biases observed when only a single visual stimulus was presented. This provided clear evidence in favour of the superposition account.

## RESULTS

### Experiment 1: Integration and recalibration biases with two visual stimuli

In the first experiment we asked how visuo-proprioceptive integration and recalibration in a cursor-control task are affected by the presence of two visual stimuli. Twenty participants performed out-and-back hand movements while receiving visual feedback about the hand endpoint (i.e., the endpoint of the outward movement) on a monitor (see Figure 1A). Specifically, proprioceptive information on hand movement endpoint (i.e., the most outward position) was accompanied by two visual stimuli (i.e., clouds of 100 small dots) that were shown briefly (100 ms) and presented with opposite spatial offsets relative to the hand position. One visual stimulus was presented synchronously with the hand reaching the movement endpoint (*Vsynch*; see Figure 1B), while the other was delayed by a semi-random interval of 750 ms with an 80 ms standard deviation (*Vdelay*; see Figure 1C). The opposite spatial offsets of the two visual stimuli relative to the hand endpoint were small and variable over trials for testing integration (see Figure 1D), and larger and constant over trials for evoking recalibration (see Figure 1E). Thus, the two visual stimuli were equated in their spatial proximity to the hand, but differed in their temporal proximity. The participants’ task was to judge the position of the hand endpoint or the position of one of the visual stimuli (these were coloured to allow their identification); which judgment to provide (hand, synchronous or delayed visual stimulus) was cued only after all stimuli of a trial had been presented. Participants responded by placing a single white dot in the visual workspace on the judged location.

**Figure. 1:**
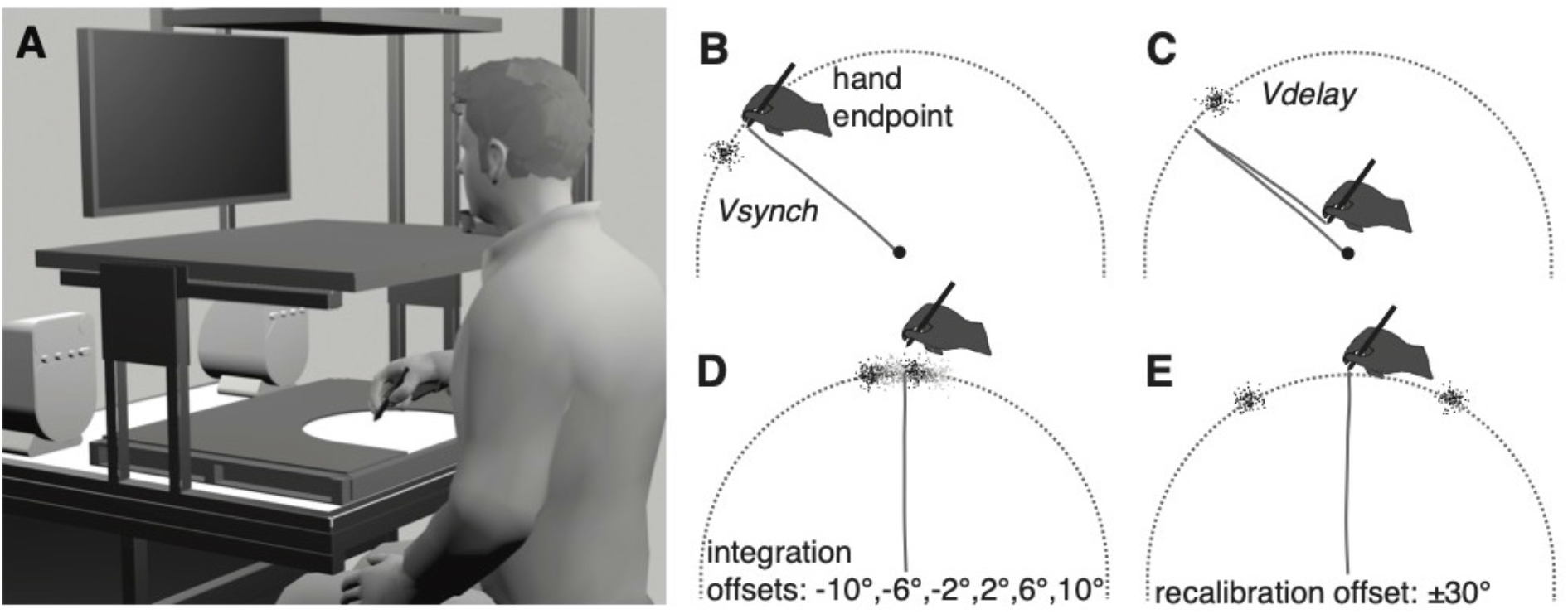
Apparatus, task, and stimuli. A) Participants performed hand movements on a digitizer tablet while visual stimuli were presented on a monitor in front of them. A semi-circular workspace was defined in the horizontal plane by a physical boundary, and in the frontal plane by an invisible semi-circle. In panels B-E, the two planes are shown on top of each other. B) When the outward-movement ended at the workspace boundary the synchronous visual stimulus was presented (*Vsynch*). Here a visual stimulus with a +10° angular offset is indicated. C) After the start of the backward-movement towards the workspace’s centre, the delayed visual stimulus (*Vdelay*) was presented around 750 ms later with an angular offset opposite to that of the synchronous stimulus (here: −10°). D) In the *bimodal judgment* trials used to assess sensory integration, the two visual stimuli had an opposite spatial offset whose magnitude varied from trial to trial (±10°, ±6°, or ±2°). E) In the *bimodal exposure* trials used to induce sensory recalibration, the two visual stimuli had opposite spatial offsets of 30° that were constant over trials. Recalibration was assessed in *unimodal judgment* trials, in which only one stimulus was present (hand, *Vsynch*, or *Vdelay*).

Our main hypotheses concerned the proprioceptive biases: we expected that the judged position of the hand endpoint would be biased towards the synchronous visual stimulus in bimodal trials with all three stimuli present (reflecting multisensory integration bias) and in unimodal trials following repeated bimodal exposure to all three stimuli (reflecting sensory recalibration bias). We also assessed the biases of the judged positions of the visual stimuli, for which we had no clear hypotheses. Individual visual stimuli are usually judged towards the position of the hand (e.g., Rand and Heuer 2013, 2019; Debats et al. 2017b), but with two visual stimuli the estimated position of each of them may be affected by both the position of the hand and the other visual stimulus.

#### Integration biases

For each participant, the biases of the judged positions of the hand, *Vsynch*, and *Vdelay* were obtained from regression analyses based on 60 *bimodal judgment* trials each, with the respective slope quantifying the integration bias (see Figure 2A). Figure 2B shows the integration biases relative to the physical stimulus locations while Figure 2C shows them such that positive values correspond to the hypothesized directions: the proprioceptive bias towards *Vsynch* and visual biases towards the hand position. Concerning our main hypothesis, we found that judgments of the hand endpoint were significantly biased towards *Vsynch* (*t*(19) = 2.88, Cohen’s d_z_ = 0.644, *p* = 0.010). This bias amounted to 12.2% of the spatial offset and pointed away from the position of *Vdelay*. In contrast, judgments of *Vsynch* were not significantly biased (*t*(19) = 0.88, d_z_ = 0.197, *p* = 0.390), while judgments of *Vdelay* were biased away from the hand position, though not significantly (*t*(19) = 1.99, d_z_ = 0.445, *p* = 0.061).

**Figure 2.**
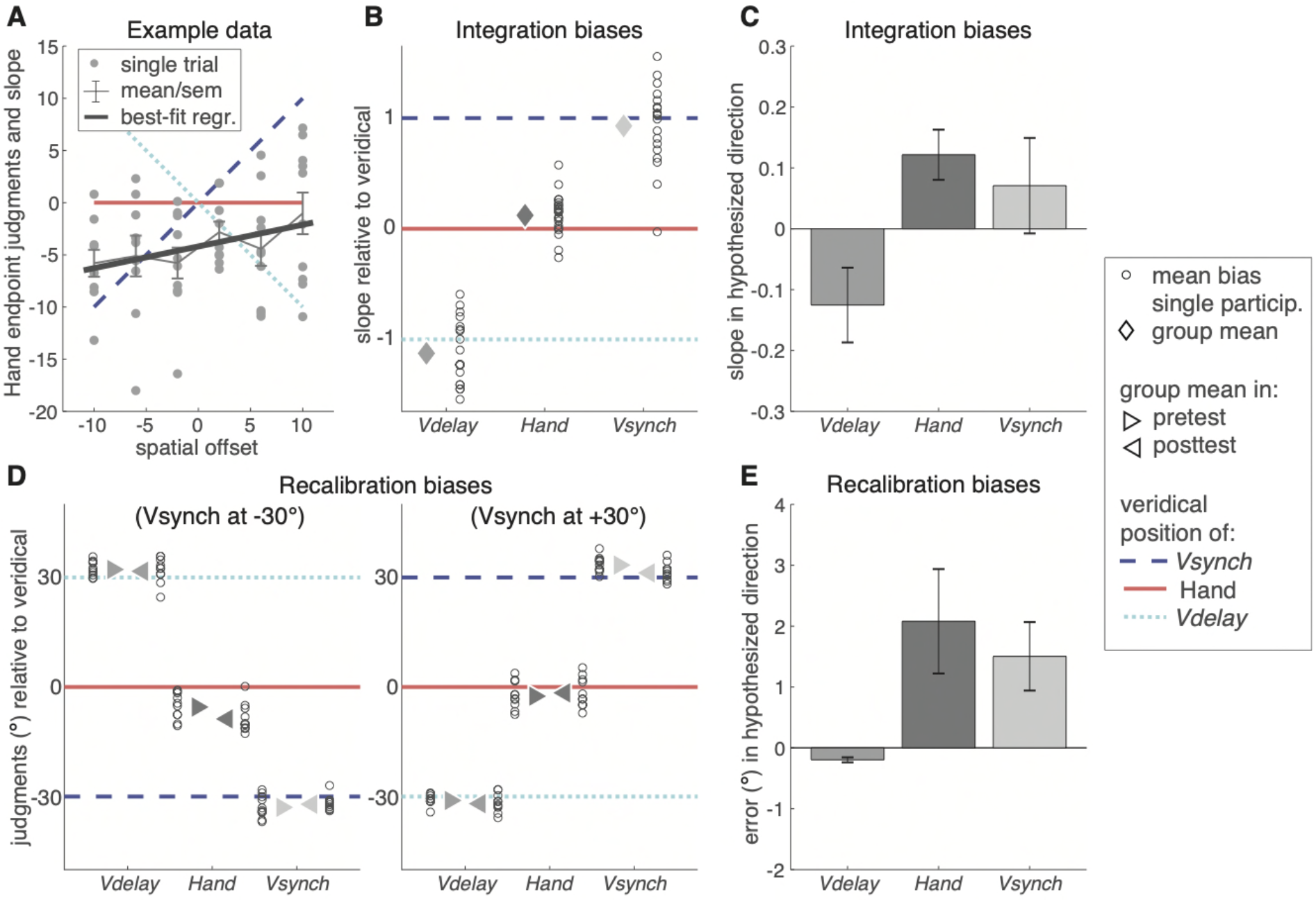
Integration and recalibration biases in experiment 1: In all panels, horizontal solid red lines indicate the veridical hand endpoint, the diagonal broken blue line that of *Vsynch*, and the diagonal dotted cyan line that of *Vdelay*. A) Integration biases were derived as slopes of linear regressions. This panel shows an example of judgments of the hand endpoints in *bimodal judgment* trials. The grey dots mark the judgment errors in individual trials. For this participant, there was a positive slope of .21, indicating that the judgments of hand position were biased towards the position of *Vsynch*. B) Individual participants’ integration biases relative to the veridical stimulus positions. The small open circles indicate individual participants’ mean slopes; diamonds indicate the group mean. C) The average integration biases (mean ± sem) in the hypothesized direction: Hand endpoint towards *Vsynch*, *Vsynch* and *Vdelay* towards the hand endpoint. D) Recalibration biases (judgment errors in degrees) for the pre-test and the post-test relative to the veridical positions of the hand, *Vsynch*, and *Vdelay*. The results are shown separately for the participants that received *Vsynch* at −30° or at +30° offset. (E) The average recalibration biases (mean ±sem) as differences between post-test and pre-test with the positive sign indicating the expected direction of the bias: hand endpoint towards where *Vsynch* had been presented in the preceding *exposure* trials, *Vsynch* and *Vdelay* towards where the hand endpoint had been in the preceding *exposure* trials.

We furthermore investigated the overall directional bias, obtained as the regression intercept. For the hand position, this was significantly different from zero and pointed in a clockwise direction (mean±sem: −3.23° ± 0.53°; *t*(19) = 5.90, d_z_ = 1.320, *p* < 0.001), whereas there were no significant overall biases for *Vsynch* (−0.42° ± 0.25°; *t*(19) = 1.65, d_z_ = 0.369, *p* = 0.116) or *Vdelay* (−0.53° ± 0.29°; *t*(19)= 1.85, d_z_ = 0.413, *p* = 0.080). In a control analysis, we quantified the intra-individual judgment variability as the standard deviation of the residuals. This variability differed between judgment types (*F*(2,38) = 7.71, ε = 0.96, *η*^2^ = 0.289, *p* = 0.002); it was largest for *Vsynch* (7.22° ± 0.29°) and smaller for the positions of *Vdelay* (6.41° ± 0.29°) or the hand (6.45° ± 0.32°). We also scrutinized the durations of the different within-trial intervals and found no differences between trials with different cued judgments (hand endpoint, *Vsynch*, and *Vdelay*) that could point to potential confounding effects of temporal delays. The duration from the start signal until the presentation of *Vsynch* varied only between 1.42 and 1.43 s between trial types (*F*(2,38) = 0.17, ε = 0.96, 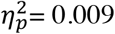, *p* = 0.842), the delay between the start signal and *Vdelay* varied between 2.48 and 2.50 s (*F*(2,38) = 0.39, ε = 0.98, 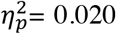, *p* = 0.682). Finally, the delay between the start signal and the judgment instruction varied between 3.02 and 3.04 s (*F*(2,38) = 0.33, ε = 0.88, 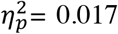, *p* = 0.723), and the delay between judgment instruction and participants’ response between 6.23 and 6.25 s (*F*(2,38) = 0.005, ε = 0.99, 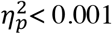, *p* = 0.995).

#### Recalibration biases

Figure 2D illustrates the biases in the *unimodal judgment* trials of pre-test and post-test relative to the stimulus positions in the *bimodal exposure* trials, separately for participants with spatial offsets of *Vsynch* of −30° and +30°, respectively. Figure 2E shows the recalibration biases such that positive values reflect a bias in the expected direction. Concerning our main hypothesis, we found that proprioceptive recalibration (bias in the judged hand position) was significant towards *Vsynch* (*t*(19) = 2.36, d_z_ = 0.529, *p* = 0.029) and thus away from *Vdelay*. There was also evidence of visual recalibration, as judgments of *Vsynch* were shifted toward the hand position in the preceding *bimodal exposure* trials (*t*(19) = 2.61, d_z_ = 0.583, *p* = 0.017), while judgments of *Vdelay* did not differ between pre- and post-test (*t*(19) = 0.38, d_z_ = 0.085, *p* = 0.708).

In addition to recalibration, we also inspected the data for motor adaptation which often accompanies proprioceptive recalibration (e.g., Hay and Pick 1966; Vijayakumar et al. 2011). It results in a shift of the direction of hand movements which partially compensates for the angular offset between cursor and hand (e.g., Krakauer et al. 2000; Bock et al. 2003). Generally, motor adaptation is stronger than sensory recalibration, and across participants the correlation between sensory recalibration and motor adaptation is small (e.g., Zbib et al. 2016; Rand and Heuer 2019). For the present data we analysed the adaptive shift of movement direction from pre-test to post-test, which (partially) compensates the offset of *Vsynch* in the preceding *bimodal exposure* trials. Motor adaptation was small and not significant (1.46° ± 1.34°; *t*(19) = 1.09, d_z_ = 0.244, *p* = 0.288). This result is consistent with the notion that motor adaptation and proprioceptive recalibration are partially independent processes (cf. Cressman and Henriques 2009, 2010; Izawa et al. 2012; Rand and Heuer 2019).

As for integration, we also probed for an overall judgment bias independent of – and thus prior to – recalibration (i.e., during the pre-test). We observed consistent judgment errors for the hand position (−3.99° ± .85°; *t*(19) = −4.60, d_z_ = 1.028, *p* < .001; Figure 2D), but not for *Vsynch* or *Vdelay* (0.22° ± 0.90° and 0.56° ± 0.53°). Control analyses showed that the intra-individual judgment variability differed between judgment types (*F*(2,38) = 5.73, ε = 0.98, 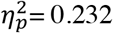, *p* = 0.007), but not between pre- and posttest (*F*(1,19) = 0.83, 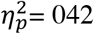, *p* = 0.375), and there was no significant interaction (*F*(2,38) = 1.76, ε = 0.97, 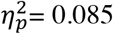, *p* = 0.187). Judgment variability was largest for the hand position (7.69° ± 0.51°) und smaller for *Vsynch* (5.99° ± 0.44°) and *Vdelay* (6.06° ± 0.43°). The durations between trial start and judgment instructions in *unimodal judgment* trials ranged between 2.8 and 2.9 s. None of the main effects was statistically significant (trial type: *F*(2,38) = 1.45, ε = 0.77, 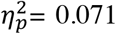, *p* = 0.248; test: *F*(1,19) = 3.05, 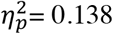, *p* = 0.097), neither was the interaction (*F*(2,38) = 1.41, ε = 0.69, 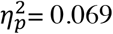, *p* = 0.256). Judgment durations were longest for *Vdelay* (7.03 ± 0.40 s), shorter for *Vsynch* (6.27 ± 0.23 s), and shortest for the hand (5.62 ± 0.17 s). These differences were statistically significant (*F*(2,38) = 8.80, ε = 0.79, 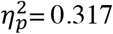, *p* = 0.002), whereas neither the difference between pre-test and post-test (6.28 ± 0.25 s versus 6.34 ± 0.22 s; *F*(1,19) = 0.06, 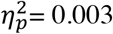, *p* = 0.808) nor the interaction (*F*(2,38) = 1.35, ε = 0.96, 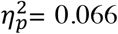, *p* = 0.271) were significant. Thus, there were no substantial differences in the within-trial timing between pre-test and post-test that could have affected the observed recalibration biases.

### Experiment 2: Winner-takes-all versus superposition hypothesis

Experiment 2 was designed to arbitrate between two alternative hypotheses on how the visual stimuli enter visual-proprioceptive combinations: i) the ‘winner-takes-all hypothesis’, according to which only the temporally most-proximal visual signal is taken into account, and ii) the ‘superposition hypothesis’, according to which integration and recalibration reflect the summed attractions by both visual signals. To contrast these hypotheses, we compared a condition in which both visual stimuli were presented (*double-stimulus* condition) with conditions in which only one of these was presented (*synchronous-stimulus* and *delayed-stimulus* conditions). We tested whether the bias of the judged hand position towards the synchronous visual stimulus was equivalent in the presence and absence of the delayed visual stimuli (in line with the winner-takes-all hypothesis); or whether the bias in the presence of two visual stimuli matched the sum of the biases induced by each of the two visual stimuli when presented alone (in line with the superposition hypothesis). For reasons of economy, we confined ourselves to the case of multisensory integration, where the within-participant comparison of the different types of trials with one or two visual stimuli is feasible. The equivalent test for recalibration would require three different participant groups who each adapt to opposite spatial offsets of two visual stimuli or to the spatial offsets of the individual stimuli.

Figure 3A illustrates the integration biases of the 22 participants for each judgment type (hand, *Vsynch*, and *Vdelay*) relative to their physical positions. Figure 3B displays the group average biases towards the hypothesized directions: positive biases for the hand position are towards *Vsynch*, while positive biases for *Vsynch* and *Vdelay* are towards the hand. With this convention the proportional bias of the judged hand position towards the position of *Vdelay* in the *delayed-stimulus* trials should be negative rather than positive, that is, away from the position of (the absent) *Vsynch*.

**Figure 3.**
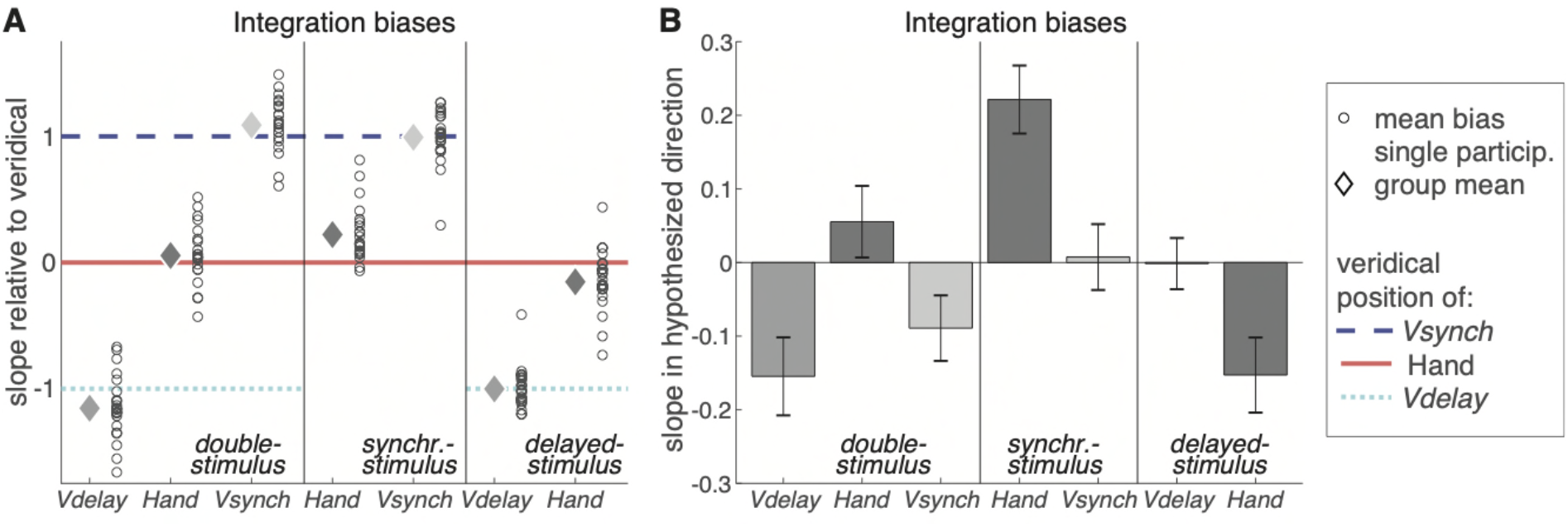
Integration bias in Experiment 2. A) Integration biases (i.e., slopes of the regression analyses) relative to the veridical positions of the hand (solid red line), of Vdelay (dotted cyan line), and of Vsynch (broken blue line) for double-stimulus, synchronous-stimulus, and delayed-stimulus trials. The open circles illustrate individual participants’ bias in the respective trials; the diamonds illustrate the group mean. B) The average integration biases (mean ± sem) in the hypothesized directions: hand endpoint towards Vsynch, Vsynch and Vdelay towards the hand endpoint. For the delayed-stimulus trials this means that the negative bias of the hand-position judgments indicates a bias away from the invisible Vsynch and towards the visible Vdelay.

In the *double-stimulus* trials, the bias of the judged hand position was positive, hence towards *Vsynch* and away from *Vdelay*. Different from Experiment 1, it was not significantly different from zero (*t*(21) = 1.11, d_z_ = 0.237, *p* = 0.278); however, the difference between experiments was not significant either (*t*(40) = 1.00, d = 0.309, *p* = 0.323), and when the data from both experiments were combined, the bias of the judged hand position was significant (0.087 ± 0.033; *t*(41) = 2.64, d_z_ = 0.407, *p* = 0.012). In *synchronous-stimulus* trials, the bias of the judged hand position towards *Vsynch* was significant (*t*(21) = 4.68, d_z_ = 0.998, *p* < 0.001). In *delayed-stimulus* trials, the bias of the judged hand position towards *Vsynch* was negative and significant (*t*(21) = 2.93, d_z_ = 0.624, *p* = 0.008), hence away from the position of (the not presented) *Vsynch* and towards the position of *Vdelay*.

We then contrasted the winner-takes-all and the superposition accounts. First, under the winner-takes-all hypothesis the bias of the hand position towards *Vsynch* should be the same in *double-stimulus* and *synchronous-stimulus* trials, whereas under the superposition hypothesis it should be larger in *synchronous-stimulus* trials: the latter was the case (*t*(21) = 2.96, d_z_ = 0.632, *p* = 0.007). Second, under the winner-takes-all hypothesis the sum of the biases in *synchronous-stimulus* and *delayed-stimulus* trials should be smaller than the bias of the judged hand position towards the position of *Vsynch* in *double-stimulus* trials, whereas under the superposition hypothesis it should correspond to that bias. In our data the sum was 0.069 ± 0.051 and not significantly different from the bias in *double-stimulus* trials, which was 0.056 ± 0.050, (*t*(21) = 0.22, d_z_ = 0.047, *p* = 0.829). Thus, the data are consistent with the superposition account and at variance with the winner-takes-all account.

The visual biases for the judgments of *Vsynch* and *Vdelay* in *double-stimulus* trials were comparable to those in Experiment 1. For *Vsynch*, the bias was close to significance (*t*(21) = 1.95, d_z_ = 0.416, *p* = 0.064), did not differ significantly from Experiment 1 (*t*(40) = 1.77, d_z_ = 0.547, *p* = 0.085), and for the combined sample it did not differ from zero (−0.012 ± 0.046 (*t*(41) = 0.280, d_z_ = 0.043, *p* = 0.781). For *Vdelay*, the bias was negative and statistically significant (*t*(21) = 2.85, d_z_ = 0.607, *p* = 0.010). It did not differ significantly from Experiment 1 (*t*(40) = 0.35, d = 0.109, *p* = 0.726), and for the combined sample it was also significant (−0.141 ± 0.041; *t*(41) = 3.440, d_z_ = 0.531, *p* = 0.001), pointing away from the hand. Thus, across both experiments, there was no reliable bias for *Vsynch* and a reliable repulsion for *Vdelay*. The visual biases in the *synchronous-stimulus* and the *delayed-stimulus* trials were close to zero (cf. Figure 3): for *Vsynch* the difference between *double-stimulus* and *synchronous-stimulus* trials did not reach significance (*t*(21) = 1.84, d_z_ = 0.392, *p* = 0.080), whereas for *Vdelay* the difference between *double-stimulus* and *delayed-stimulus* trials was significant (t(21) = 2.65, d_z_ = 0.565, p = 0.015). Thus, the repulsion of *Vdelay* in *double-stimulus* trials depended on the preceding presentation of *Vsynch*, implying that the repulsion was not away from the position of the hand, but from the position of *Vsynch*.

The overall biases, as estimated by the regression intercepts, and the intra-individual variability were comparable to Experiment 1. The overall bias for hand judgments, averaged across trial types, was clockwise (−3.18 ± 0.71°), while no consistent bias emerged for the visual stimuli (*Vsynch*: −0.06 ± 0.28°; *Vdelay*: −0.12 ± 0.28°). Judgment variability in *double-stimulus* trials differed according to judgment type (*F*(2,42) = 11.17, ε = 0.92, 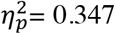, *p* < 0.001); as in Experiment 1, variability was largest for *Vsynch* (8.32° ± 0.37°) and smaller for *Vdelay* (6.81° ± 0.28°) or the hand (7.22 ± 0.35°). Also, as in Experiment 1, the analysis of the various within-trial durations revealed no systematic variations across trial types that could have affected the main findings. The duration from the signal to start the movement until presentation of *Vsynch* was 1.24 ± 0.04 s in *double-stimulus* and 1.25 ± 0.04 s in *synchronous-stimulus* trials (*t*(21) = 1.45, d_z_ = 0.309, *p* = 0.162), and until presentation of *Vdelay* it was 2.22 ± 0.05 s in *double-stimulus* and 2.23 ± 0.05 s in *delayed-stimulus trials* (*t*(21) = 0.58, d_z_ = 0.124, *p* = 0.566). The time to the judgment instruction was 2.64 ± 0.07 s, 2.67 ± 0.07 s, and 2.66 ± 0.07 s in *double-stimulus, synchronous-stimulus*, and *delayed-stimulus* trials, respectively, (*F*(2,42) = 1.48, ε = 0.81, 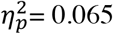, *p* = 0.242), and the judgment duration was 5.80 ± 0.21 s, 6.00 ± 0.24 s, and 5.77 ± 0.21 s in the three types of trials (*F*(2,42) = 3.10, ε = 0.95, 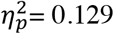, *p* = 0.058).

## DISCUSSION

Multisensory integration and sensory recalibration are two processes by which our brain exploits redundant multisensory signals. While integration serves to reduce the immediate discrepancies between two signals by merging them into a combined estimate, recalibration reduces consistent discrepancies by aligning the unisensory estimates. Importantly, for multisensory perception to deliver meaningful results in every day settings, our brain has to first identify the relevant redundant signals for integration and recalibration (Talsma et al. 2010). While previous studies on integration or recalibration biases have investigated multisensory perception based on paradigms presenting one stimulus per modality (for exceptions see: Bertelson et al. 2000; Tong et al. 2020), we here asked how two stimuli in one modality are combined with the evidence in another modality. We found that when two visual stimuli accompany a hand movement, proprioceptive judgments of the hand position are influenced by both visual signals, though the temporally more proximal visual stimulus exerts a stronger influence.

### Superposition of multiple multisensory attractions

When accompanied by two visual stimuli, the judged endpoint of a hand movement was biased towards the position of a synchronous visual stimulus and not towards the delayed visual stimulus. Similarly, after a series of exposure trials in which movements were consistently accompanied by the two visual stimuli with opposite spatial offsets, there was a recalibration bias of the judged hand endpoint towards the synchronous visual stimulus. Hence, the synchronous visual stimulus dominated both integration and recalibration over the delayed visual stimulus. This dominance could result from a winner-takes-all interaction between the two visual stimuli, where only the dominant one attracts the perceived position of the hand. Alternatively, it could result from the superposition of two biases, where the stronger attraction by the synchronous visual stimulus is superposed on the weaker attraction by the delayed one. We contrasted these two hypotheses for integration and found support in favour of the superposition hypothesis.

We propose that this superposition principle also holds for recalibration. Firstly, the proprioceptive integration bias with two visual stimuli was clearly smaller than in a condition with the synchronous visual stimulus only as in experiment 2 and as observed previously (Debats and Heuer 2018b). Similarly, the proprioceptive recalibration bias with two visual signals that we observed here, amounted to about 7% of the visuo-proprioceptive offset only and was smaller than the order of 20% typically found with single visual stimuli (Henriques and Cressman 2012; Barkley et al. 2014; Zbib et al. 2016; Rand and Heuer 2019). This suggests that both visual stimuli influenced the recalibration process in the present study. Furthermore, integration and recalibration bear many functional similarities, as both are responses to redundant multimodal signals: one is the immediate effect and the other is the respective aftereffect. Also, both are governed by the reliability rule (van Beers et al. 2002; Burge et al. 2010; Debats et al. 2017b), both are supposedly steered by perceptual causal inference (e.g., Körding et al. 2007; Sato et al. 2007; Badde et al. 2020; Debats and Heuer 2020a) and previous neuroimaging studies have provided support that similar brain regions support multisensory recalibration and integration (Park and Kayser 2019, 2021).

The finding that the temporally most proximal stimulus dominates the multi-modal perceptual judgment seems a straightforward consequence of the role of temporal proximity during integration and recalibration in general (Talsma et al. 2010). In particular, temporal proximity was shown to influence the perceived redundancy of multiple stimuli as well as judgments of their causal relation (Wallace et al. 2004; Parise et al. 2012) and is a strong driver for the cross-modal association of stimuli in general (Talsma et al. 2010). Thereby the perceived causal relation is considered a key driver of multisensory integration and sensory recalibration (Körding et al. 2007; Sato et al. 2007; Badde et al. 2020). With this in mind, the present results suggest that the stronger influence of the synchronous compared to the delayed visual stimulus reflects the perceived stronger causal relation between the synchronous visual stimulus and the hand movement. While a number of studies showed that consistent temporal relations facilitate the perceptual pop-out, for example in search tasks (Van der Burg et al. 2008, 2011; Alsius and Soto-Faraco 2011), few studies have tested this hypothesis for the perceptual biases arising from multisensory integration or recalibration.

One could raise the concern that differences other than the relative timing of the two visual stimuli in the trial sequence may have contributed to the observed results. Specifically, the difference in temporal proximity of the two visual stimuli to the hand movement was accompanied by a difference of the time to the judgment instruction at the end of the trial: the delay between the synchronous visual stimulus and the start of the judgment was about 1.5 s, whereas it was only about 0.5 s for the delayed visual stimulus. We argue that this is a minor concern for two reasons. First, a one-second difference is only small in relation to the duration of the judgment itself, which was about 6 s. Second, a recent study on audio-visual integration and recalibration has directly tested the effect of temporal delays and concluded that any potential influence of memory decay is small, at best (Park and Kayser 2020). Thus, the critical temporal offset that affects integration and recalibration should be the one between stimuli, not the one between stimuli and start of the judgment.

### Repulsion among visual stimuli

In the present data, the judgments of the visual stimuli were only weakly influenced by the proprioceptive information on the hand endpoint. Nevertheless, there was a remarkable pattern of biases in the judged positions of the delayed visual stimuli. First, for integration we found that in trials with both visual stimuli present, the perceived position of the delayed visual stimulus was consistently repulsed from the position of the hand and thus also from the position of the synchronous visual stimulus. In contrast, this repulsion of the delayed visual stimulus was absent in trials without the synchronous visual stimulus (Experiment 2). This suggests that the delayed visual stimulus was repulsed by the synchronous visual stimulus rather than by the proprioceptive signals. Second, for recalibration we observed no repulsion of the delayed visual stimulus away from where the synchronous visual stimulus had been presented during the exposure trials. Hence, repulsion of the judged position of the delayed visual stimulus was present only in trials in which its presentation had been preceded by the synchronous visual stimulus.

We envisage two potential explanations for this repulsion. First, the spatial attraction between the perceived positions of the hand at the end of a movement and its visual consequence has a parallel in the temporal attraction between a simple action (e.g. keypress) and its sensory effect – a phenomenon known as ‘intentional binding’ (Haggard et al. 2002b; Moore and Obhi 2012; Kirsch et al. 2016). Such binding has been observed both with auditory stimuli following or preceding a keypress, while for two successive auditory stimuli or keypresses attraction was absent or turned into repulsion (Haggard et al. 2002a). Hence, mutual attraction not only in time, but also in space, may be limited to cross-modal stimuli (proprioceptive and visual in the present experiments) and may not exist for intra-modal pairs (two distinct visual stimuli in the present experiments). Second, at the time the delayed visual stimulus was presented, participants may still have attended, or even fixated, the position of the synchronous visual stimulus. Thus, the distance of the delayed visual stimulus from the synchronous one (and therefore also the hand position) may have been overestimated because of its retinal eccentricity (Bock 1986, 1993; Henriques and Crawford 2002). Alternatively, the perceived position of the delayed stimulus could have been subject to an attentional repulsion effect (Pratt and Turk-Browne 2003; Pratt and Arnott 2008; Baumeler et al. 2020). Although our finding of a repulsion between the perceived positions of the visual stimuli had not been expected and its explanation is rather tentative, it demonstrates that the mutual relations between the perceived positions of an ensemble of multimodal stimuli are shaped by different influences, not only by superposition of mutual attractions.

### Absence of visual integration bias

Different from similar previous experiments (Debats et al. 2017a, b; Debats and Heuer 2018a, 2020b, a), particularly one with a highly comparable condition (Debats and Heuer 2018b, condition “End”), we observed no reliable integration biases for the judged positions of the visual stimuli when only one of them was presented. Although visual integration biases were generally small in related previous studies, their absence was unexpected. We propose that this relates to two unique features that distinguish experiment 2 from these previous studies. First, the visual stimuli were clouds of dispersed dots presented for 100 ms only, whereas previously solid circles had been used which were presented for longer durations. Besides making the estimates of the visual positions comparatively unreliable, this appearance of the visual stimuli was also quite different from the appearance of a cursor in every day computer use and could have reduced the stimuli’s presumed redundancy. Second, in Experiment 2, trials with a single visual stimulus were randomly mixed with trials with two visual stimuli during the out-and-back hand movements. These double-stimulus trials might also have corrupted the presumed redundancy of the cross-modal stimuli because a single position of the hand cannot correspond to two different positions of visual stimuli. Overall the presumed redundancy could thus have been smaller than in those previous studies, resulting in a reduced integration strength that would explain the small and unreliable visual integration bias observed in experiment 2.

### Limitations and outlook

We set out to study multisensory perception in settings with more than one pair of redundant cross-modal stimuli, a step towards more natural situations where redundant signals for integration and recalibration have to be identified among a plethora of signals (Nardo et al. 2014; Spence and Frings 2020). We directly confirmed that the use of redundant stimuli for multisensory integration follows a superposition principle. We propose that this principle also holds for recalibration, a hypothesis to be tested in future studies employing more time efficient paradigms. Similarly, future work should extend these results to other combinations of sensory modalities, such as audio-visual ventriloquist paradigms (e.g., Park and Kayser 2020).

## ACKNOWLEDGEMENTS

This study was funded by the Deutsche Forschungsgemeinschaft (DFG KA2661/2-1) We thank A. Sonntag, P. Unterbrink, and S. Böker, for their support in data collection.

## COMPLIANCE WITH ETHICAL STANDARDS

The authors declare that no conflicts of interest exist. All procedures were approved by the Bielefeld University Ethics Committee and in accordance with the 1964 Helsinki declaration. Informed consent was obtained from all individual participants included in the study.

## AUTHOR CONTRIBUTIONS

Designed the experiments: NBD, HH, CK; Analysed the data: HH; Wrote the manuscript: NBD, HH, CK

## DATA AVAILABILITY STATEMENT

All data analyzed in this study are available at https://pub.uni-bielefeld.de/record/2954727:

## METHODS

### Participants

We collected data from 20 participants for Experiment 1 (aged 18 to 41 years, mean ± sd: 25.3 ± 5.9 years; 15 female) and 22 participants for Experiment 2 (aged 19 to 30 years; mean ± sd: 24.5 ± 2.9 years; 10 female). According to G*Power 3.1.9.2 (Faul et al. 2007), this sample-size is sufficient to detect medium-sized integration and recalibration effects with an error probability of 0.05 and power of 0.8. (Effect size d_z_ was set to 0.6, which is about half the effect size computed from the integration biases found by Debats and Heuer (2018b) and less than half the recalibration biases reported by Rand and Heuer (2019) for experimental paradigms similar to the present one.) In Experiment 1, the data of five additional participants was replaced because after data screening only few valid trials remained (see below). All participants were right handed according to a German version of the Edinburgh handedness inventory (Oldfield 1971). The experiments were conducted in accordance with the declaration of Helsinki and approved by the Bielefeld University Ethics Committee. Participants gave written informed consent prior to participation and were compensated with a payment of €7 per hour.

### Apparatus

Participants sat in front of a table with their head stabilized by a chinrest. A digitizer tablet (Wacom Intuos4 XL; 48.8 by 30.5 cm) was placed on the table, and a computer monitor (Samsung MD230; 23 inches; 50.9 by 28.6 cm) was mounted at 60 cm viewing distance (Figure 1A). Participants held a stylus with their right hand as during writing and pressed a button on it with their thumb or index finger to submit responses. The stylus could be moved on the digitizer within a semi-circular workspace of 15 cm radius that was bounded by a 5 mm thick PVC template. The boundary served as a mechanical stop for outward movements. A horizontal opaque board prevented the direct vision of hand and stylus. The position of the stylus was recorded at 60 Hz with a spatial resolution of 0.01 mm, using a custom-built MATLAB program with the Psychophysics Toolbox (Kleiner et al. 2007). During the experiment the room lights were off. All stimuli on the monitor were presented in light grey on a black background with the exceptions described below.

### Task

A general task-description is provided here, more detail on task and stimuli are given below and in Figure 4. Participants made out-and-back movement with their unseen right hand on the digitizer tablet, with each movement proceeding from the centre of the semi-circular workspace to its boundary and back to the centre. Most crucial was the endpoint of the outward movement (i.e., where the hand touched the boundary). To prevent stereotypy in these endpoints, an approximate direction for the hand movement was cued, centred at one of five possible angles (60°, 75°, 90° (straight ahead), 105°, or 120). Two visual stimuli on the monitor (clouds of dots) could provide visual information on hand endpoint. One visual stimulus appeared once the hand reached the boundary (the synchronous visual stimulus: *Vsynch*; Figure 1B), and the other was delayed such that it appeared during the backward movement (the delayed visual stimulus: *Vdelay*; Figure 1C). The visual stimuli had an opposite spatial offset from the hand endpoint. The offset is necessary to reveal integration and to induce recalibration.

**Figure 4:**
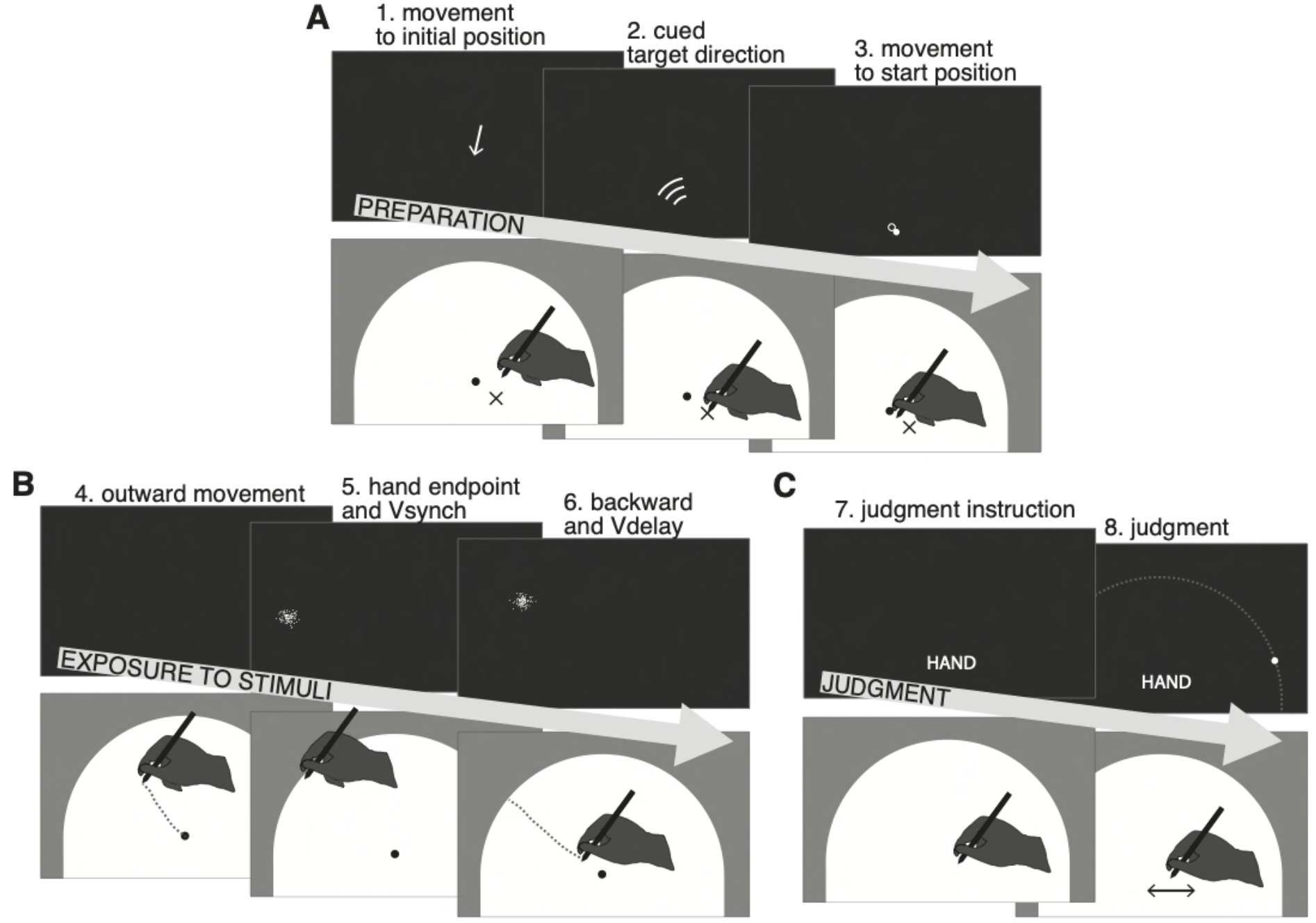
Experimental trials. A) During the preparation phase of each trial, participants were guided by arrows on the monitor to an initial position, and a direction for the outward movement was cued. Participants then moved the cursor to the start position in the centre of the workspace. B) During the out-and-back movement, different stimuli could be presented. Depending on trial type, this could be the *hand endpoint* (i.e., the position of the tip of the stylus when hitting the workspace boundary), and/or the visual stimulus (cloud of dots) synchronous with the hand reaching the workspace boundary (*Vsynch*), and/or the delayed visual stimulus, presented around 750 ms after starting the backward movement (*Vdelay*). One of the visual stimuli was blue, the other yellow. Both were located on the invisible semi-circle that corresponded to the workspace boundary. C) In trials that included a position judgment, a written instruction was shown on the monitor to indicate which position was to be judged: the hand endpoint, the (centre of the) blue cloud, or the (centre of the) yellow cloud. Judgments were indicated on the monitor by means of steering a small dot to the desired position on the semi-circular track that corresponded to the workspace boundary. (The track is shown here for reasons of clarity, but was not visible to the participants.)

After completion of the backward movement, participants were cued to indicate either the perceived hand endpoint or the position of one of the two visual stimuli. They did so by moving a small white dot on the monitor along an invisible semi-circular track that corresponded to the workspace boundary. Judgments of hand endpoints were thus cross-modal (proprioceptive to visual) and required a reference-frame transformation (horizontal plane to fronto-parallel plane), whereas judgments of the positions of the visual stimuli were intra-modal and required no such transformation. According to previous work (Debats and Heuer 2018a), the different frames of reference have only marginal or no effects. Yet cross-modal judgments are noisier than intra-modal judgments and therefore affect the asymmetry of the biases, leading to a relative enhancement of the biases for the estimated hand positions in the present experiments (Debats et al. 2017b).

### Visual stimuli

Visual stimuli were clouds of 100 dots of 1 mm diameter each. Their positions had a bivariate normal distribution with a standard deviation (SD) of 5 mm. Dots that deviated by more than ±3 SD from the centre on one of the coordinates were moved into this range by subtracting the deviation from the position at 6 SD from the centre. The resulting clouds of dots thus had a diameter of 30 mm (corresponding to 2.86° of visual angle, or 11.46° in the polar coordinates with the origin in the centre of the workspace). The bivariate normal distribution was centred on the polar coordinates (α+ρ, r), with r the radius of the invisible semi-circle (150 mm), α the angular position of the movement endpoint, and ρ the angular spatial offset between visual stimulus and movement endpoint (positive ρ was counter clockwise, negative ρ was clockwise). For testing integration, we used small and randomly varying angular spatial offsets (−10°, −6°, −2°, 2°, 6°, 10°; Figure 1D); for inducing recalibration we used larger consistent offsets (+30° and −30°; Figure 1E).

The synchronous visual stimulus was presented for 100 ms when the stylus passed a threshold of 97% of the workspace radius from the centre. This 97%criterion was just below the movement distance at which the stylus would hit the boundary. The delayed stimulus was presented for 100 ms with a normally distributed delay (mean: 750 ms, SD: 80 ms) after the backward movement had passed the 97% threshold.

Visual stimuli were either presented in blue (RGB: 0 155 155) or yellow (RGB: 155 155 0); for half of the participants the synchronous stimulus was blue and the delayed one was yellow, for the other half this was reversed. The different colours were used to facilitate judgment instruction (‘blue cloud’ or ‘yellow cloud’). The colours assigned to *Vsynch* and *Vdelay* were counterbalanced over the participants.

### Trial types

Experiment 1 comprised three types of trials that differed with respect to the stimuli presented (hand endpoint, *Vsynch*, *Vdelay*) and the judgment required (hand endpoint, *Vsynch*, *Vdelay*, none). In *bimodal judgment trials*, all three stimuli were presented and the position of one of them had to be judged. In *bimodal exposure* trials, all three stimuli were presented, but no judgment was required. In *unimodal judgment* trials, only one out of the three stimuli was presented and its position judged. Thus, only a movement was performed or a ‘synchronous’ or ‘delayed’ visual stimulus was presented. When a visual stimulus was presented without a movement being performed, the terms ‘synchronous’ and ‘delayed’ refer to shorter or longer delays after the trial onset; these were adopted from the timing of *Vsynch* and *Vdelay* in an immediately preceding *bimodal exposure* trial. Similarly, the judgment instruction was presented at a similar time compared to the preceding trial, and the spatial offset was defined relative to an assumed hand endpoint that corresponded to the movement error in the preceding trial plus the currently cued movement direction (which was cued although no movement was to be performed).

Experiment 2 comprised three variants of *bimodal judgment* trials. *Double-stimulus* trials were identical to the *bimodal judgment* trials of Experiment 1. In *synchronous-stimulus* trials only *Vsynch* was presented when the outward movement reached the workspace boundary, and in *delayed-stimulus* trials only *Vdelay* was presented during the backward movement.

### Experimental design

Experiment 1 comprised an initial practice block of 20 trials, a first part to assess integration, and a second part to assess recalibration. First, integration was tested in 180 *bimodal judgment* trials. These were 2 repetitions of 90 unique trials: 3 types of position judgments (hand endpoint, *Vsynch, Vdelay*), 6 pairs of spatial offsets of *Vsynch* and *Vdelay* (−10°/10°, −6°/6°, −2°/2°, 2°/−2°, 6°/−6°, 10°/−10°), 5 cued ranges of target directions (60°, 75°, 90°, 105°, or 120°). The 90 trials were presented in two blocks of 90 trials each, the trial order being randomized within each block.

Second, recalibration was tested in a series of 300 trials. Initially there was a pre-test of 60 trials, 30 *bimodal exposure* trials alternating with 30 *unimodal judgment* trials (10 hand, 10 *Vsynch*, and 10 *Vdelay* in semi-randomized order). It was followed by the exposure phase of 180 *bimodal exposure* trials that varied only in the cued target direction. Half of the participants were exposed to *Vsynch* and *Vdelay* at +30° and −30°, for the other half this was reversed. Finally, the post-test of 60 trials was identical to the pre-test except for the unique randomized trial order. Trials were arranged in three blocks of 90 trials (pre-test and exposure), 120 trials (exposure), and 90 trials (exposure and post-test), which were of approximately equal durations.

Experiment 2 comprised an initial practice block of 20 trials followed by 420 experimental trials. These were 2 repetitions of 210 unique trials: 90 were *double-stimulus* trials, 60 *synchronous-stimulus* trials, and 60 *delayed-stimulus* trials. These 90 or 60 trials varied in the 3 or 2 types of position judgments (hand endpoint, *Vsynch*, and/or *Vdelay*), the 6 pairs of spatial offsets of *Vsynch* and *Vdelay* (−10°/10°, −6°/6°, −2°/2°, 2°/−2°, 6°/−6°, 10°/−10°), and the 5 cued ranges of target directions (60°, 75°, 90°, 105°, or 120°). The 420 trials were presented in six blocks of 70 trials each, trial order being randomized within each set of 210 trials.

### Detailed trial description

Trials were composed of two or three phases: preparation, exposure to the stimuli, and judgment. In the preparation phase (Figure 4A) an arrow on the monitor directed participants to an initial starting position which was randomly selected within a rectangular area just below (−20 to −40 mm) and to both sides (−15 to +15 mm) of the centre of the semi-circular workspace. One second after reaching the initial position, a WiFi-like symbol was presented for one second. It consisted of three 15° arcs at 18, 24, and 30 mm radial distance from the centre of the workspace and cued one of the five potential ranges of directions for the outward-movement. These symbols were also presented in *unimodal judgment* trials in which no movements were performed because they also indicated a range of directions in which visual stimuli were likely to appear. Then, the start position in the centre of the workspace was marked by an outline circle (7 mm diameter) and the position of the hand relative to the start position was indicated by a filled circle (6 mm diameter). Once the start position was reached and the stylus remained for 500 ms within a tolerance range of 2.5 mm from its centre, the outline circle and the filled circle disappeared and an auditory beep of 0.3 s duration was presented as a go-signal for the outward movement. The hand movement from the initial position to the start position was included to force participants’ eyes off a target location for their outward movement. The colour of the outline circle indicated the current type of trial. It was light grey in all bimodal trials (i.e., *bimodal judgment* and *bimodal exposure*). In *unimodal judgment* trials without movement it was red, instructing participants to stay at the start position, and in *unimodal judgment* trials with movement it was green.

The preparation phase of each trial was followed by the exposure phase (Figure 4B). The beginning of the outward movement was recorded when the distance from the start position exceeded 2.5 mm, and its end was defined when 97% of the 150 mm distance to the workspace boundary were passed. The movement reversal was defined as the time period from passing the 97% threshold in the outward direction until passing that threshold in the backward direction. The backward movement started when this threshold was passed and ended (signalled by a short beep) when the position of the hand had not changed by more than 2.5 mm for 500 ms. In *bimodal judgment* and *bimodal exposure* trials, *Vsynch* and *Vdelay* were presented at the appropriate times, and in *unimodal judgment* trials only a movement was performed or only a visual stimulus was presented with its timing and position as described above.

In the judgment phase (Figure 4C) of *bimodal* and *unimodal judgment* trials, the type of judgment was instructed upon completion of the backward movement and after the visual stimuli had been presented. (In *unimodal judgment* trials without a movement the timing was adopted from the preceding *bimodal exposure* trial.) A short text was presented on the monitor below the start position, ‘hand’, ‘yellow cloud’ or ‘blue cloud’, and remained visible until the end of the judgment. One second after the instruction, a light grey filled circular marker (2 mm diameter) appeared at a random position on the invisible semi-circle that corresponded to the workspace boundary. Participants could navigate the marker along the semi-circular path by way of small movements of the stylus to the left and right relative to its position at the end of the backward movement (or the start position when no movement had been performed). When the position of the marker matched the remembered hand endpoint or the remembered position of the instructed visual stimulus, whichever had to be judged, participants pressed the button on the stylus to confirm their judgment. There were no time constraints for this.

### Data pre-processing

The primary data for the analysis of integration and recalibration were the judgment errors. As in our previous studies (e.g., Debats and Heuer 2018b), in addition to screening the judgment errors for outliers, we screened the movement trajectories for deviations from the instructions which required smooth outward movements in the cued direction followed by immediate backward movements. The following criteria were used (in brackets the percentages of discarded trials are indicated separately for judgment trials and exposure trials): (1) the absolute angular deviation between the physical and judged positions of a visual stimulus or the hand endpoint was larger than 20° (judgment trials Exp.1, Exp. 2: 3.29%, 5.88%), (2) the outward movement included a reversal of more than 10 mm (judgment trials Exp.1, Exp. 2: 0.69%, 0.45%; exposure trials Exp. 1: 1.17%). (3) the stylus slid along the workspace boundary for more than 2.5° during movement reversal (judgment trials Exp.1, Exp. 2: 3.23%, 3.41%; exposure trials Exp. 1: 5.17%), (4) the movement direction deviated more than 30° from the centre of the instructed range of movement directions (judgment trials Exp. 1, Exp. 2: 2.13%, 3.17%; exposure trials Exp. 1: 2.54%), and (5) the hand moved away from the start position by more than 15 mm in trials in which no movement was to be performed (judgment trials Exp. 1: 0.46%). For Experiment 1, the medians of the number of included trials per participant were (ranges are in brackets): *bimodal judgment* trials with hand position judged: 56.5 (35-60), with *Vsynch* position judged: 55.5 (41-60), and with *Vdelay* position judged: 56.5 (34-60); *unimodal judgment* trials with hand position judged: 17.5 (8-20), with *Vsynch* position judged: 19.0 (17-20), and with *Vdelay* position judged: 19.0 (18-20). The five participants who were replaced by new participants had more than 60% discarded trials in at least one of the different types of judgment trials; three of them made many movements that did not obey the instructions, and two of them made many large errors in judging the hand position. For Experiment 2, the median (and the minimum-maximum) of the number of retained trials for the 7 different types per participant were: *double-stimulus* trials with judgments of hand endpoint: 53.5 (33-59), of *Vsynch*: 50.0 (29-60), and of *Vdelay*: 54.0 (37-59); *synchronous-stimulus* trials with judgments of hand endpoint: 53.0 (35-59) and of *Vsynch*: 53.0 (35-58); *delayed-stimulus* trials with judgments of hand endpoint: 54.0 (38-59) and of *Vdelay*: 56.0 (39-60).

### Dependent variables to quantify integration

Our primary interest was i) the proprioceptive bias, that is, the bias of the judged hand position towards the position of *Vsynch*. As the spatial offsets of *Vsynch* and *Vdelay* were symmetric around the hand endpoint, any bias of the hand position towards *Vsynch* was also a bias away from *Vdelay*. In addition, we quantified ii) the visual biases of the judged positions of both visual stimuli towards the position of the hand, iii) the consistent judgment errors that were independent of the spatial offsets between visual stimuli and hand position, and iv) the intra-individual variability of the judgments. As the timing of the events in each trial depended on the precise movement kinematics, we also quantified v) the duration of individual phases of the experimental trials, such as between reaching the start position and the presentation of *Vsynch* or *Vdelay*, or the duration of the judgments.

Biases, consistent judgment errors, and judgment variability were determined from regression analyses performed for each participant and judgment type separately. Specifically, for each *bimodal judgment* trial we computed the judgment error, that is, the deviation of the judged from the physical angular position of the hand or a visual stimulus (positive errors were counter clockwise, negative errors were clockwise). These judgment errors were regressed against the angular spatial offsets, i.e. the difference between the physical angular positions of visual stimulus and hand endpoint. The slope of this regression provides a measure of the proportional bias, that is, of the judgment error as a proportion of the angular offset. The intercept reflects consistent judgment errors independent of the visual stimuli, and the standard deviation of the residuals can serve as measure of intra-individual judgment variability. Figure 2A shows an example for an individual participant’s judgments of hand endpoints.

### Dependent variables to quantify recalibration

Recalibration was quantified based on the *unimodal judgment* trials during pre-test and post-test. We report the mean judgment errors during the pre-test and the recalibration bias. The latter was computed as difference between the judgments in post-test and pretest with the sign of the difference chosen such that positive differences are adaptive: for hand judgments a positive bias represents a shift towards the position of *Vsynch* in the preceding *bimodal exposure* trials; for visual judgments a positive bias represents a shift towards the hand position in the preceding *bimodal exposure* trials. Motor adaptation was assessed as the change of the movement directions in *unimodal judgment* trials from pre-test to post-test. Positive values are adaptive: they represent a shift of the movement direction away from the position of *Vsynch* in the preceding *bimodal exposure* trials.

### Statistical analyses

For both experiments we tested the regression slopes and intercepts against zero using two-sided t-tests. We further compared the biases between Experiments 1 and 2 using independent-samples t-tests and tested them against zero for the combined sample. For recalibration the adaptive biases in judgments and movement directions were tested against zero using t-tests as well. For the main purpose of Experiment 2, the following two analyses were implemented. First, we compared the proportional biases of the judged hand position towards *Vsynch* in *double-stimulus* and *synchronous-stimulus* trials. Second, the proportional bias of the judged hand position towards *Vsynch* in *doublestimulus* trials was compared with the sum of the biases in *synchronous-stimulus* and *delayed-stimulus* trials. As effect size for these tests we report d_z_, which is the ratio of mean and standard deviation of the measurements (for paired-sample t-tests the mean and standard deviation of the differences are entered).

Judgment variability was compared using an ANOVA with the within-participant factors judgment type and pre-test vs post-test (for recalibration). Corresponding ANOVAs served to analyse the durations of various trial phases. ANOVA results are reported with Greenhouse-Geisser epsilon and adjusted *p*. As effect size we report partial eta-squared, 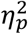, the ratio of the sum of squares for an effect divided by the sum of the sums of squares for the effect and the error.

## Notes

https://pub.uni-bielefeld.de/record/2954727

### Competing Interest Statement

The authors have declared no competing interest.

